# LOICA: Logical Operators for Integrated Cell Algorithms

**DOI:** 10.1101/2021.09.21.460548

**Authors:** Gonzalo Vidal, Carlos Vidal-Céspedes, Timothy James Rudge

## Abstract

Mathematical and computational modeling is essential to genetic design automation and for the synthetic biology design-build-test-learn cycle. The construction and analysis of models is enabled by abstraction based on a hierarchy of components, devices, and systems that can be used to compose genetic circuits. These abstract elements must be parameterized from data derived from relevant experiments, and these experiments related to the part composition of the abstract components of the circuits measured. Here we present LOICA (Logical Operators for Integrated Cell Algorithms), a Python package for modeling and characterizing genetic circuits based on a simple object-oriented design abstraction. LOICA uses classes to represent different biological and experimental components, which generate models through their interactions. High-level designs are linked to their part composition via SynBioHub. Furthermore, LOICA communicates with Flapjack, a data management and analysis tool, to link to experimental data, enabling abstracted elements to characterize themselves.

## 2 INTRODUCTION

Synthetic Biology is an interdisciplinary field that mixes life sciences and engineering. From this perspective living systems are objects to engineer, and a rational way to design them is by modifying their genetic code. This can be done by introducing synthetic DNA that encodes a synthetic regulatory network, also known as a genetic circuit. The design-build-test-learn (DBTL) cycle is central to engineering disciplines and each phase requires appropriate tools, standards and workflows, which are still in development in synthetic biology. The synthetic biology open language (SBOL) is an open standard for the representation of *in silico* biological designs that covers the DBTL cycle and has attracted a community of developers that have produced an ecosystem of software tools [4].

Modeling is key to the DBTL cycle and is essential to the design and learn stages; a model states a well-defined hypothesis about the system operation. Abstraction enables the construction and analysis of models based on components, devices, and systems that can be used to compose genetic circuits. It is the basis for genetic design automation (GDA), which can accelerate and automate the genetic circuit design process. In order for GDA to proceed in a rational way, the abstract elements of genetic circuits must be accessible to characterization, allowing parameterization of models of their operation and interactions.

Functional abstraction of DNA sequences as parts such as transcriptional promoters (Pro), ribosome binding sites (RBS), coding sequences (CDS), terminators (Ter) and other elements has enabled the assembly of relatively small genetic circuits [1–3]. However, for large-scale genetic circuit design higher-level abstractions are required, as provided by the logic formalism [6]. In this approach circuit compositions are abstracted into genetic logic gates that transition between discrete low and high steady-state gene expression levels according to input signals, either external or internal to the circuit [9]. These genetic logic circuits can be designed automatically, in an analogous way to electronic circuits, based on the required discrete logical truth table [6], however this specification requires knowledge of the domain-specific programming language *Verilog*.

Despite the discrete logical design formalism, these genetic circuits are dynamical systems and can have autonomous, continuous non-steady-state dynamics, displaying complex and rich behaviors from bi-stability to oscillations and even chaos [2, 3, 11]. Furthermore, typical operating conditions for engineered circuits like colonies, bioreactors or gut microbiomes are time varying, which can lead to complex behaviors from even simple genetic circuits [7].

To design genetic circuit temporal dynamics we therefore require kinetic gene expression data generated at the test phase. This data must be integrated with models to enable characterization of abstracted parts, devices and systems, as well as metadata, including the DNA part composition, to enable automated design. Thus there is a need for software design tools that integrate abstract circuit designs, dynamical models, kinetic gene expression data, and DNA part composition via common exchange standards in a user-friendly and accessible fashion.

## 3 RESULTS

Logical Operators for Integrated Cell Algorithms (LOICA) provides a high-level genetic design abstraction using a simple and flexible object-oriented programming approach in Python. LOICA integrates models with experimental data via two-way communication with Flapjack, a data management and analysis tool for genetic circuit characterization [12]. This communication not only provides direct access to experimental characterization data, but also to DNA design composition and sequence via SBOL contained in SynBioHub [5], enabling characterization and simulation in the same tool, but also facilitating exchange with other tools such as iBioSim [11].

The basic objects in LOICA are *Operator* and *GeneProduct*, which may be either a *Regulator* or *Reporter* (Figure 1A). A *Regulator* represents a molecular species that regulates gene expression. A *Reporter* is a molecular species that provides a measureable signal, such as a fluorescent protein. The *Operator* maps one or more *Regulator* concentrations to one or more *GeneProduct* synthesis rates. An *Operator* can be implemented in DNA as a combination of Pro and RBS, and the *Regulator* could be a CDS of a transcription factor or of a regulatory RNA. The interactions between the *Operators* and the *Regulators* encode models for genetic circuit temporal dynamics, which are simulated with differential equations. The system is thus:

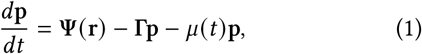

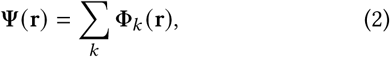

where (**p** = *p*_0_, *p*_1_, …*p*_*N*−1_)^*T*^ is the vector of *GeneProducts*, which includes different *Regulators* (**r** = (*r*_0_, *r*_1_, …*r*_*M*−1_) ^*T*^) and *Reporters* (**s** = (*s*_0_, *s*_1_, …*s*_*N* − *M*−1_)^*T*^). The non-linear operator **Ψ** maps *Regulator* concentrations to *GeneProduct* synthesis rates. **Γ** is a diagonal matrix of *GeneProduct* degradation rates, and *μ*(t) is the instantaneous growth rate of the cells. Equation 1 shows the overall system where **Ψ** encodes the whole circuit, and consists of a sum of individual LOICA *Operators* **Φ**_*k*_ (Equation 2).

**Figure 1:**
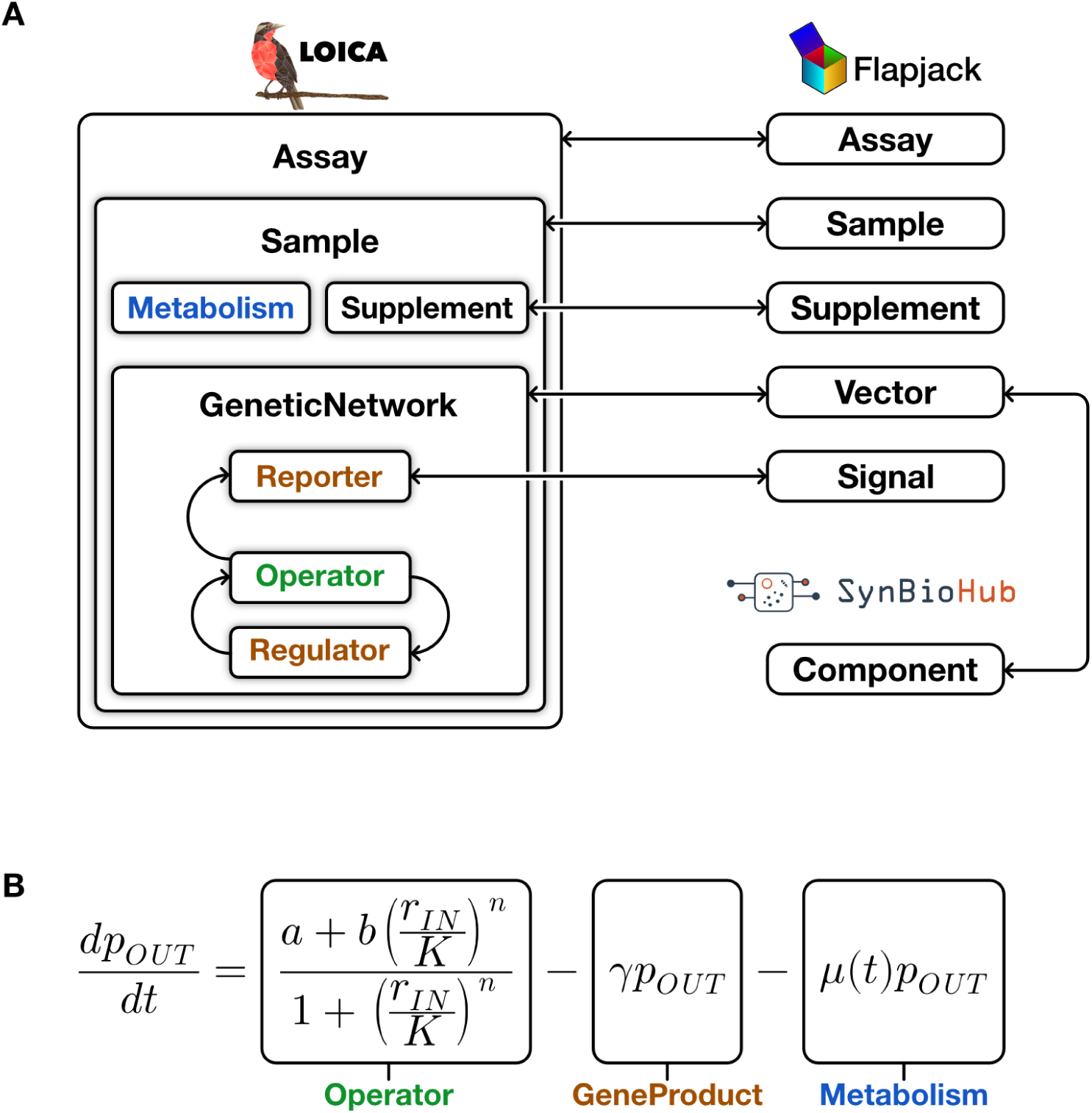
Diagram of model generation in LOICA. A. Diagram of an *Assay* encapsulating a *Sample* which in turn encapsulates *Metabolism, Supplement*, and *GeneticNetwork*. In the later, the *Operator* and *Regulator* are interacting to generate a model. On the right side the different interactions with the Flapjack and SynBioHub models are shown. B. Mathematical model of a NOT *Operator* with one input and one output generated through LOICA object interactions. Here *p*_*OUT*_ is a *GeneProduct* (*Regulator* or *Reporter*), output of the *Operator*. In the *Operator a* is the basal or leaky gene expression, *b* is the regulated gene expression, *r*_*IN*_ is a *Regulator* concentration, *K* is the switching concentration, and *n* is the cooperativity degree of *r*_*IN*_ with respect to the *Operator*. In the *GeneProduct γ* is the degradation rate of *p*_*OUT*_. In *Metabolism μ* (*t*) is the instantaneous growth rate which dilutes *p*_*OUT*_. Here the *Operator* is encoded by a Hill equation transfer function that states its regulation by *r*_*IN*_. The transfer function could be any mapping from input concentration to synthetis rate, making LOICA *Operators* flexible and easy to extend.

Figure 1B shows a mathematical model that results from the interaction of an *Operator* encoding a simple NOT logic modeled with a Hill equation as transfer function. It can be implemented as a promoter containing repressor binding sites combined with a ribosome binding site (RBS). The logical *Operator* can thus be instantiated as a genetic device that is repressed by an input *Regulator* and outputs a *GeneProduct* synthesis rate. Note that LOICA can be used to define an *Operator* as any operation that maps from input *Regulator* concentrations to output synthesis rates.

The repressilator is a useful dynamical system case study because it produces continuous sustained oscillations that escapes ON/OFF logic [2]. To model it, we consider a simple balanced ring oscillator with three NOT *Operators* connected with three different *Regulators*. The *Operators* and *Regulators* are incorporated into a *GenticNetwork*, linked with a Flapjack *Vector*, which with the *Metabolism* drives the dynamics of the *Sample*, also corresponding to a Flapjack *Sample* (Figure 1A). For the circuit to produce a measurable signal we add three *Operators* using the same inputs but changing the outputs to three different *Reporters*, linked with the Flapjack *Signal* model (Figure 2A). The code to generate this model is in Figure 2B. This approach is used to generate synthetic data (a LOICA *Assay*) from models that can be uploaded to Flapjack. It is then easy to access Flapjack’s genetic circuit characterization tools, data management and data visualization through a Python package (pyFlapjack) or the web interface shown in Figure 2D.

**Figure 2:**
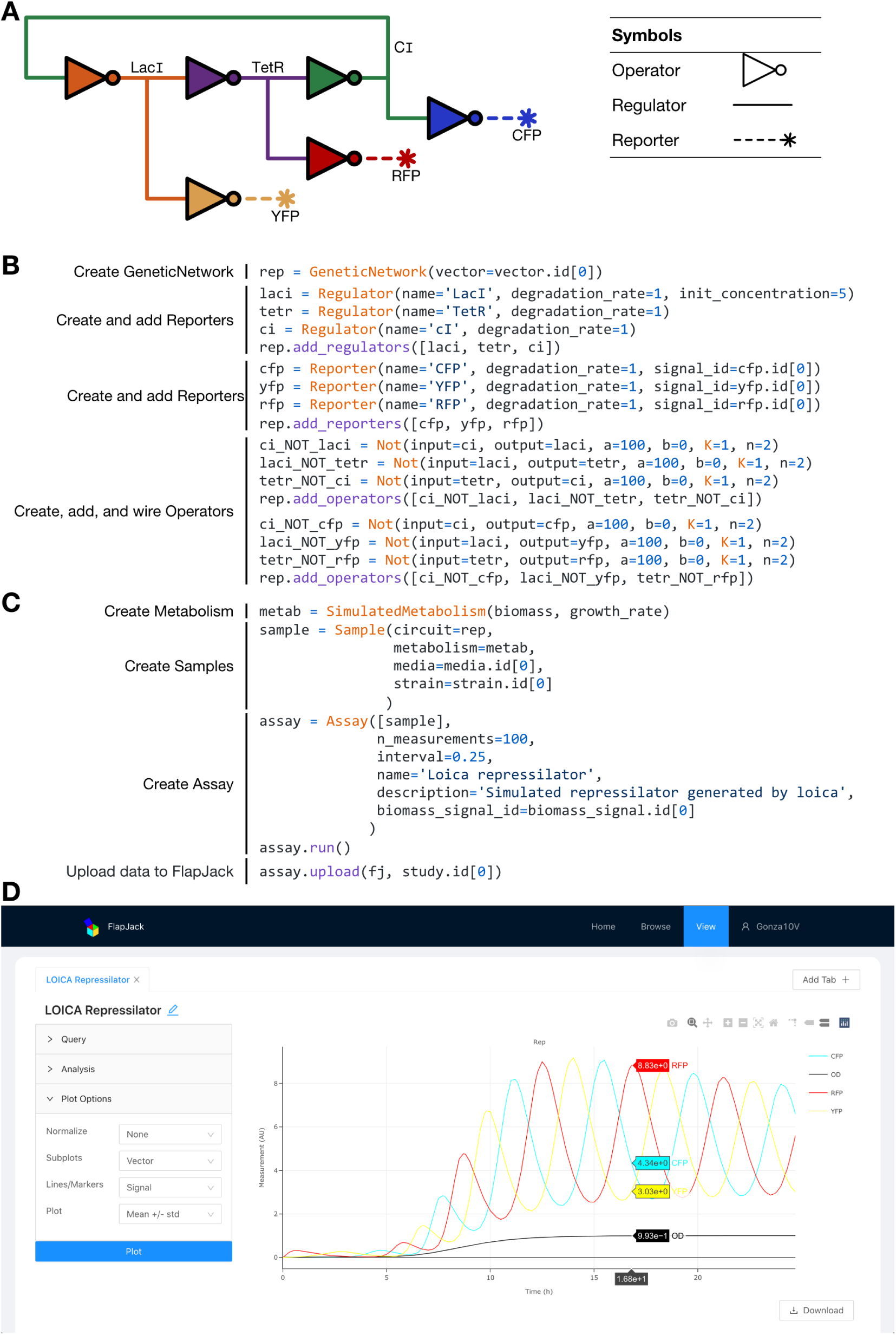
Example of repressilator design in LOICA. A. LOICA diagram of the modeled repressilator circuit and respective symbols. B. Python code that generates a repressilator in LOICA. GeneticNetwork construction is the first step where the user states all the objects and their interaction. C. Next during *Assay* setup the user initializes and runs the simulation, and the results can be uploaded to Flapjack. The two-way communication with Flapjack allows data storage and management, enables various analyses to be performed, and allows *Operators* to characterize themselves. D. Data exploration in Flapjack via the web interface. This interface allows to query, analyze and plot the data, which can also be done through the pyFlapjack Python package.

We have described how to use LOICA to generate and analyze simulation data from models. Another example workflow goes from data to model parameterization and could be as follows for the learn stage of the DBTL cycle. First, a *GeneticNetwork* is assembled from a collection of *Operators*, linked by various *Regulators* and *Reporters* in some topology. Each *Operator* is then linked to experimental data contained in Flapjack, which corresponds to measurements of the auxiliary circuits required to parameterize the model encoded. For example, a NOT *Operator* links to data of a chemical signal receiver and a chemical signal inverter measured in a range of signal concentrations (LOICA and Flapjack *Supplement*). The *Operator* provides a function that then extracts the data from Flapjack and uses it to parameterize the corresponding model (Figure 1B). In the example shown that means fitting parameters *a, b, K, n* by least squares minimization of the difference between the experimental data and the solution of differential equation models of the auxiliary circuits (Equation 1). Each *Operator* thus contains the information required to characterize itself. Therefore with LOICA data driven models that include mathematical constraints it is possible to design circuit topologies and look for likely functional designs initiating a new round of the DBTL cycle.

## 4 CONCLUSION

LOICA integrates the design and characterization of genetic circuit dynamics into Python workspaces, providing an easy-to-understand design abstraction implemented using simple object-oriented programming principles. This programming interface does not require specialist or domain-specific knowledge, but leverages common programming skills, making it accessible but also providing customization capabilities for advanced users. LOICA is able to simulate abstract genetic circuit designs using differential equation models. It abstracts genetic circuit designs into objects which are capable of characterizing themselves via links to data in Flapjack. As data relating to genetic components is updated in Flapjack, the fitted parameters can be automatically updated upon characterization. Flapjack also provides the connection to SynBioHub which allows design and characterization based on SBOL.

## 5 FUTURE WORK

We aim to complete and automate the DBTL cycle in synthetic biology, proposing a workflow that integrates LOICA, Flapjack [12] and SynBioHub [5] via SBOL3 [4]. In the design stage SBOL/SBML will provide flexibility to use other existing bioCAD tools such as iBioSim [11] as part of the workflow. To make a direct transition to the build stage LOICA will generate a human/machine readable protocols for assembly using Opentrons liquid handling robots [10] and their open Python API. We will expand LOICA’s capabilities to single-cell resolution spatio-temporal systems by connecting it to CellModeller [8] for individual based modeling and stochastic simulations. For the test stage we will develop open science hardware for measuring genetic circuit dynamics that could be used to obtain kinetic data for direct upload to Flapjack.

